# Identification of chemo-sensory genes from antennae of female banana pseudostem weevil, *Odoiporous longicollis* by transcriptomics approach

**DOI:** 10.1101/2022.06.16.496379

**Authors:** Kannan Mani, Padmanaban Balakrishnan, Victoria Soroker

## Abstract

In the present study, the female *O. longicollis* antennal transcriptome was constructed by Illumina Hiseq 2500™ sequencing, with the aim of discovering olfaction-related genes. Totally 12, 411 unigenes were identified from the transcriptome assembly and the putative genes functions were annotated using gene ontology tools. We identified 46 putative chemosensory unigenes likely essential for insect olfaction including 19 odorant binding proteins, 3 Niemann–Pick type C (NPC) protein, 6 odorant receptors, 15 ionotropic receptors, and 3 sensory neuron membrane proteins. The key function of these chemosensory genes in the antennae are discussed. Phylogenetic analysis of NPC revealed that the identified proteins had a close relationship with Coleopterans than other insects. This is the first ever report on identification of olfactory genes from *O. longicollis* which may provide new leads for control of this major pest.

## Introduction

Insect chemo-sensing organ is located in the antennae that are used to detect food sources, oviposition sites, aggregation, prey avoidance and finding sexual partner for their survival and reproduction (Hildebrand and Shepherd 1997; Li *et al*. 2014; 2015). Insects receive the volatiles from environment, it is carried by odorant binding proteins to olfactory receptor neurons (ORNs) where they transform the odour information into neuronal electrical signals (Leal 2013). Many genes are reported to be participate in the molecular network of olfaction such as Niemann-Pick disease, type C1 and 2 (NPC1-2), Odorant binding protein (OBP), olfactory receptors (ORs), Ionotropic receptors (IRs), Sensory Neuron Membrane Protein (SNMP) (Vogt 2003; Pelosi *et al*. 2006). To date, various studies are reporting the presence of chemo-olfactory related genes from antennae of many coleopteran insect species (*Dendroctonus ponderosae, Ips typographus, Rhynchophorus palmarum* and *Cylas formicarius*) but their genome have not been sequenced yet (Andersson *et al*. 2013; Bin *et al*. 2017; Gonzalez et al., 2021). However, Next-generation RNA sequencing is an advanced technique along with bioinformatic tools provides a huge genetic information’s of non-model organism without any prior sequence knowledge (Liu *et al*. 2015). Therefore, studies on the identification of chemosensory genes in insects are making more significant progress and it could be used to succeed in gene silencing mediated environmentally friendly pest management (Leal *et al*. 2008; Nganso *et al*. 2021).

The banana pseudostem weevil (BPW), *Odoiporus longicollis* (Coleoptera: Curculinoidae) is a monophagous pest of *Musa* sp. The banana pseudostem weevil is a serious pest and threat to banana production because it causes crop losses from 10 to 90 % depending on the management efficiency. The *O. longicollis* widely found and distributed throughout South-East Asian countries (Padmanaban *et al*. 2020). The cryptic feeding behavior of grub and nocturnal activities of adult weevil making more difficult to control this pest using chemical pesticides. In addition, the use of pesticides shows insect resistance and leads to off target effects on natural enemies and other living animals (Prasuna *et al*. 2008). Eco-friendly approaches like clean planting material, crop sanitation, monitoring weevil in the garden using longitudinal split banana stem trap, longitudinal split banana stem trap with color light emitting diode, use of entomopathogenic fungi as biocontrol agents and botanical pesticide are still followed by farmers but unable to completely eradicate the pest (Kannan *et al*. 2021; Kannan *et al*. 2021). Pheromone (2-methyl-4-heptanol) with host plant extract mixture used as trap showed significant attraction of stem weevil (Palanisamy *et al*. 2019). However, there is no report on molecular mechanism of olfaction in *O. longicollis*.

A preliminary information has been reported on BPW chemosensory organ using scanning electron microscopy, consist of basal scape, pedicel, and a flagellum chemoreceptive sensillae in the antenna. The SEM studies indicated that each segment of antennal club bears three types of sensory organs such as the sensilla type I, II and III (Nahif *et al*. 2003). The sensilla are involved in recognizing suitable host for oviposition when many cultivars are present in the same vicinity. Electroantennogram studies indicated considerable differences among the host plant volatiles (Prasuna *et al*. 2008; Alagesan *et al*. 2018). Screening of *Musa* germplasm against banana pseudostem weevil also indicated similar difference in feeding (Padmanaban *et al*. 2001). These researches are clearly indicating the importance of BPW antennae on chemo-olfaction (**Fig.1**). Therefore, a detailed study on identification of chemosensory genes from female antennae of banana pseudostem weevil is prerequisite because the specific groups of genes play a major role in detecting host plant for feeding and oviposition preference. In this context, the present study was focused on to identify the genes from antennae responsible for olfaction of female *O. longicollis* using Whole transcriptome sequencing.

**Fig. 1.**
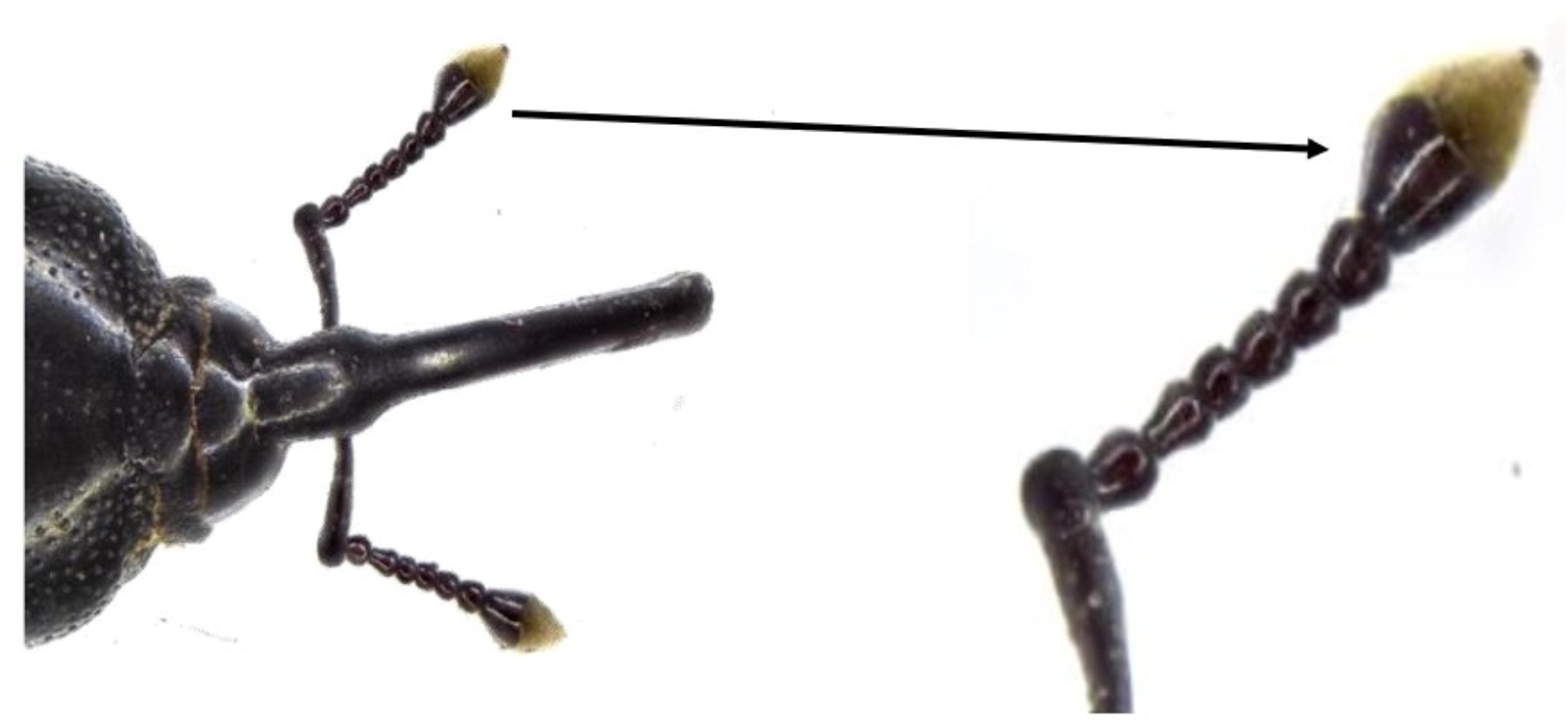
Morphology of Antennae of banana pseudostem weevil, *O. longicollis*

## Material and methods

### Sample collection

BPWs were collected from field and maintained in the laboratory as described by Padmanaban *et al*. (2020). The newly emerged weevil fed with fresh pieces of pseudostem. The ten days old female weevils of *O. longicollis* were used for collection of antennae. The antennae were dissected (150 nos of females) using sterile surgical blade (Size: 11) by immobilizing the insect on ice and collected antennae into 1.5mL Eppendorf tube containing RNAlater® solution (Thermofisher, USA). Fifty pairs of weevil antennae were used per replicate. The antenna samples were transported to Xcelris Labs Limited, Ahmedabad-380015, Gujarat, India in dry ice for the further analysis.

### RNA isolation, Illumina 2x 150 paired end library preparation and sequencing

The total RNA was extracted from the antennae by TRIzol method (Invitrogen, USA). The quality and quantity of the isolated RNA was checked by 1% formaldehyde denaturing Agarose gel and Qubit ® 2.0 flurometer respectively. The RNA was purified from genomic DNA contamination by TURBO DNA-free™ Kit following the manufacturer’s protocol (Thermo Fisher Scientific, Lithuania, USA). Libraries were prepared from purified RNA (∼1 μg) using Illumina TruSeq standard mRNA library preparation kit as per manufacturer protocol (Illumina Inc., San Diego, CA). Briefly, total RNA was subjected to Oligo dT beads to enrich mRNA fragments, then subjected to purification, fragmentation and priming for cDNA synthesis. The fragmented mRNA was converted into first-strand cDNA, followed by second-strand cDNA synthesis, A-tailing, adapter-index ligation and finally amplified by recommended number of PCR cycles. The amplified libraries were analyzed on Bioanalyzer 2100 (Agilent technologies) using High sensitivity DNA chip as per manufacturer’s instructions and sequenced on the Illumina Hiseq 4000 platform.

### *De novo* transcriptome assembly and gene annotation

Clean reads were obtained by removing reads containing adaptor sequences, more than 5% unknown nucleotides, more than 50% bases with Q-value ≤ 20 and empty reads (Trimmomatic, version: 0.36). Then, the Q30 and GC-content were used to assess the sequencing quality. After filtration of raw data, master assembly (*de novo*) was performed from triplicates using Trinity software (Ver. 2.1.1) with default assembly parameters, kmer 25 to generate unigenes. All unigenes generated from transcript, CDS prediction and functional annotation against NR and TF database using CD-hit (ver. 4.6.1), Transcoder (Ver. 2.0.1) and Blastp (Ver. 2.2.30) respectively. All the assembled unigenes were searched against the Nr (NCBI non-redundant protein sequences), Nt (NCBI nucleotide), KOG (eukaryotic ortholog groups), Swiss Prot, KEGG (Kyoto Encyclopedia of Genes and Genomes) database using BLASTx and BLASTn with an E-value < 10^−5^. Blast2 GO was used for GO (Gene Ontology) annotation with an E-value < 10^−5^ based on the protein annotation results of the Nr database. Inter Pro functional annotation was performed using InterProScan7.

### Chemosensory gene identification, alignment, and phylogenetic analysis

The chemosensory related genes were identified from the annotated name of the BLASTx hits, or based on the GO terms (molecular function, biological process, and cellular component) as detected by the blast2go analysis (keywords: odorant binding, ionotropic receptor, NPC2, SNMP, response to pheromone). The female banana stem weevil antennal chemosensory transcripts open reading frames (ORF) were translated into protein for phylogenic analysis. The NPC proteins sequences of *O. longicollis* were analyzed using an HMM profile to obtain the closely related proteins sequences of different organisms. The alignment was used to search the ‘reference proteomes’ database (non-redundant proteome sets) for significant matching proteins using HMM search [e-value < e-5, HMMER web version 2.21.2 (Finn *et al*. 2015) with a more permissive significance threshold (e-value< 0.00001).

In this study, four beetles *(Tribolium castaneum, Dendroctonus ponderosae, Asbolus verrucosus, Agrilus planipennis)* and one non beetle organism *(Drosophila melanogaster)* as out group were selected for classification, annotation, and phylogenetic study. The NPC genes from each organism and *O. longicollis* sequences were aligned by MAFFT in “Auto” strategy method with BLOSUM62 scoring matrix and other default parameters (Katoh and Standley 2013). NPC gene sequences from each organism were realigned, and conserved blocks were subjected to a maximum likelihood analysis using PhyMl in 500 bootstraps (Guindon *et al*. 2010). For each analysis, the best model was selected by PhyMl SMS (Lefort *et al*. 2017). Branch support was estimated using a fast and accurate maximum likelihood-ratio test (aLRT) and subsequently the visualization and graphically edit of the circular phylogenetic tree were conducted using the iTOL server (Letunic and Bork 2016).

## Results and Discussion

### Illumina sequencing

In the present study, we intended to identify the chemo-sensory genes from antenna of female *O. longicollis* using the Illumina Hiseq™ 2500 platform as a step toward understanding olfactory machineries in this weevil. The total raw reads of 29191558, 38305324 and 24594454 for sample R1, R2 and R3 respectively were obtained from cDNA libraries. After filtration of raw data produced 19328370, 25685810 and 16993976 clean reads for R1, R2 and R3 respectively. These reads were assembled into 12411 unigenes (accumulated length of 13763296 bp) with a mean unigene length of 1108 bp and N50 of 1754 bp. The unigenes size distribution analysis indicated that the lengths of the 3872 (31.19 %) were greater than 1000 bp **(Fig. 2**).

**Fig. 2.**
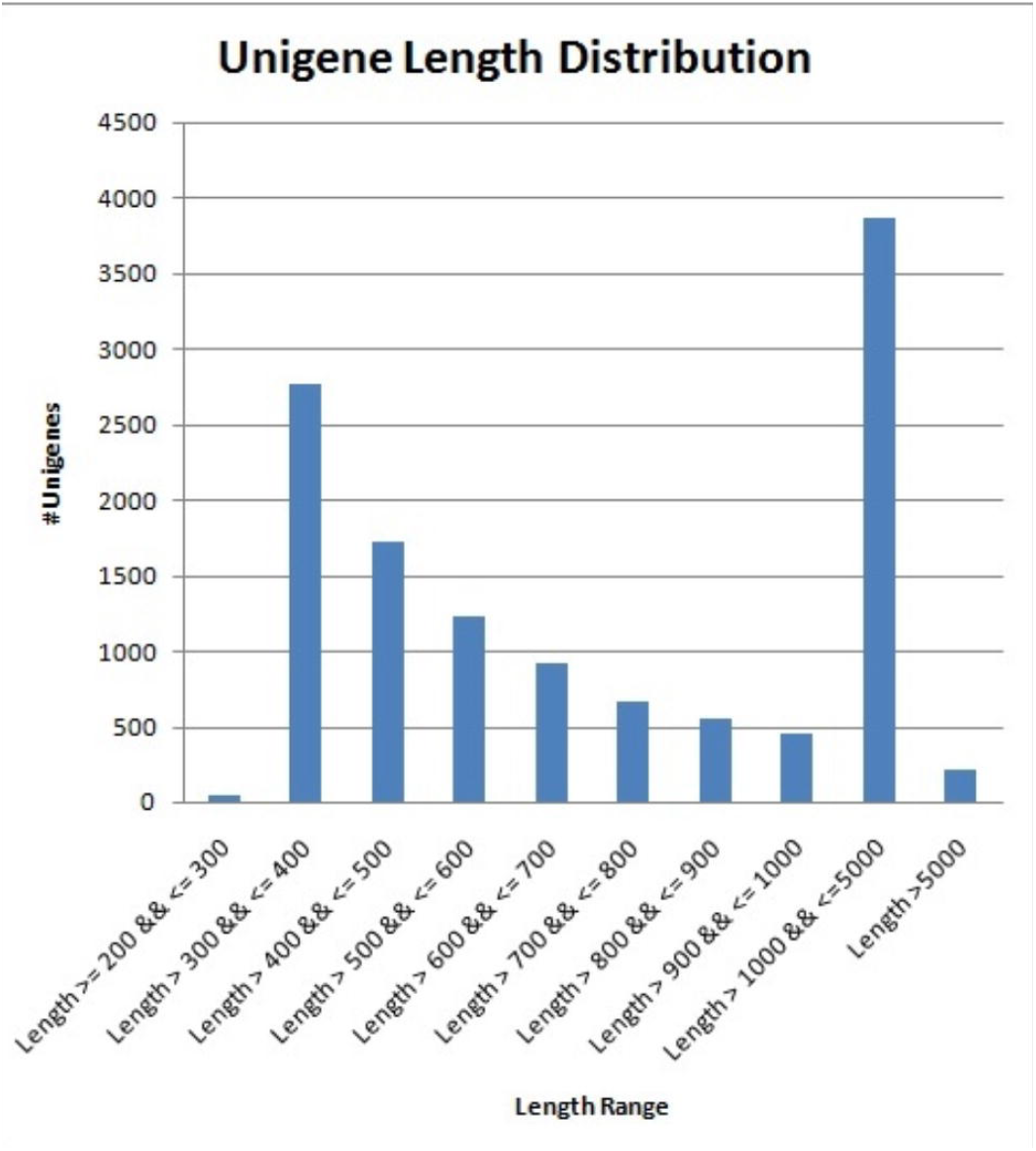
Unigenes size distribution.

### Functional annotation of assembled unigenes

In the transcriptome sets, we identified 12, 411 unigenes with a mean length of 1108 bp and, a maximum length 26, 640 bp, it was less when compared with previous studies (Bin *et al*. 2017). Annotation was conducted at least one of the databases by BLASTx and BLASTn program with the E-value cut-off of 10^−5^ for about 12,411 unigenes. 10, 199 (82.17 %) unigenes were annotated by the NCBI-Nr database, 8260 (66.55 %) unigenes by Uniprot, 2711 (51.94 %) by GO, 6447 (39.07 %) by the COG database and 4850 (21.84 %) by Pfam database. In molecular function, the most abundant transcripts in the antennae were linked to binding and catalytic activity. In biological process, the most represented biological processes were cellular, metabolic, and biological regulation. In the cellular component terms, membrane and cell, and organelle constituted the most abundant categories. These GO assignments are in accordance with previous antennal transcriptomics of coleopteran insects (Hu *et al*. 2016; Wu *et al*. 2016; Antony *et al*. 2016; Wang *et al*. 2016, Liu *et al*. 2015) (**Fig. 3A**). The above-mentioned functions are involved in the part of the chemo-sensing mechanism (Leal 2013; Pelosi *et al*. 2006). Similarly euKaryotic of Orthologous Genes (KOG) analysis clearly shows that most of the genes (gene count:1372) are highly involved signal transduction mechanism (**Fig. 3B**). This result indeed shows the significant contribution of antennal proteins in chemo-olfaction of *O. longicollis*. BLASTx homology searches in the NCBI-Nr database showed that *O. longicollis* antennal transcriptomes had a best blast match to coleopteran sequences, primarily the mountain pine beetle *Dendroctonus ponderosae*, (80%) and *Anoplohora glabripennis* (68%) and less to *Leptinotarsa decemlineata* (48%) and the red flour beetle *T. castaneum* (42%) which indicates that *O. longicollis* evolutionarily conserved among coleopteran insects.

**Fig. 3.**
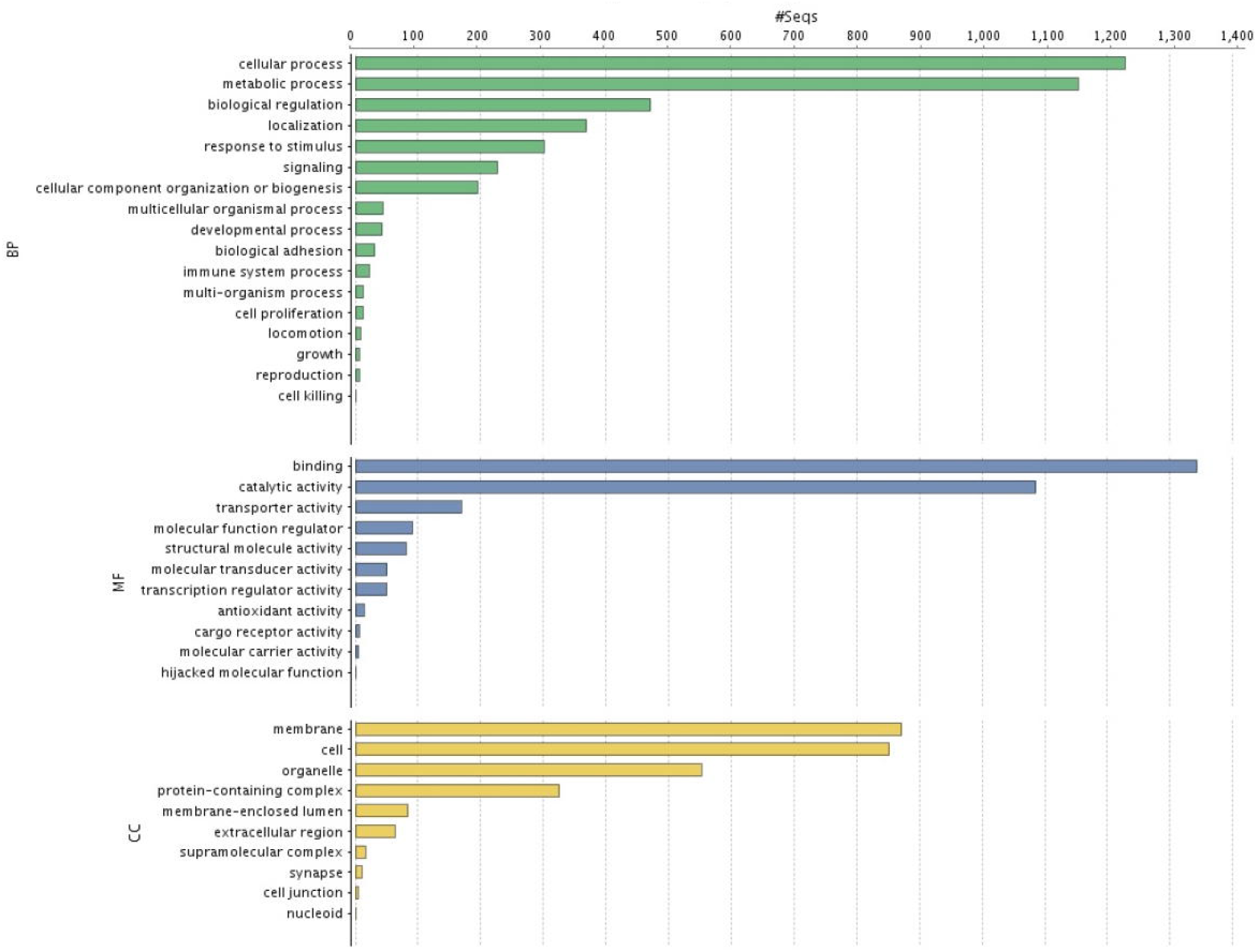
**A** Gene ontology (GO) classification of unigenes obtained female antennal transcriptome and were classified into 3 functional categories: molecular function (MF), biological process (BP), and cellular component (CC). **Fig. 3.B** Classification of unigenes based on Clusters of Orthologous Groups (COG)

### Gene families associated with odorant detection

Gene families involved in odorant detection such as odorant binding proteins (OBPs: 19), odorant receptors (ORs: 6), ionotropic receptors (IRs:15), and three sensory neuron membrane proteins (SNMPs) were identified in female *O. longicollis* antennal transcriptomes (Table. 1). Information of candidate genes (ORs, IRs, SNMPs and OBPs) such as the gene name, unigenes sequences, lengths, predicted protein sequences and the annotation in NCBI-Nr database, predicted protein domains are listed in the Supplementary file 1. These genes are not only involved in chemo sensing, but also in metabolic activity. Low number of transcripts were identified from the present study when compared with antennal transcriptome of *A. glabripennis* where it has 40 OBPs, 12 CSPs, 37 ORs, 11 GRs, 4 IRs, 2 SNMPs and 1 ODE (Hu *et al*. 2016). The difference between the organism may be due to less expression of those genes or species specific (Wang *et al*. 2017).

### Identification of NPC family

We identified one transcript of NPC1 and two transcripts of NPC2 were detected according to the presence of the conserved domain (myeloid differentiation (MD)-2-related lipid-recognition, IPR003172). The NPC1 transcript found in *O. longicollis* clustered with *D. ponderosae* with highly significant bootstrap values (**Fig. 4**). Niemann-Pick proteins of class 1 and 2 are act as carriers for cholesterol and lipids. Recently, NPC considered as a third class of binding proteins for semio-chemicals in arthropods because of their binding affinity for small hydrophobic compounds (Pelosi *et al*. 2014). The banana stem and spadix contain saponin, phytosterol, sterols and steroids (Ismail *et al*. 2018; Choudhury *et al*. 2019) are may be act as attractant and those compounds may be carried by NPC proteins for host selection.

**Fig. 4.**
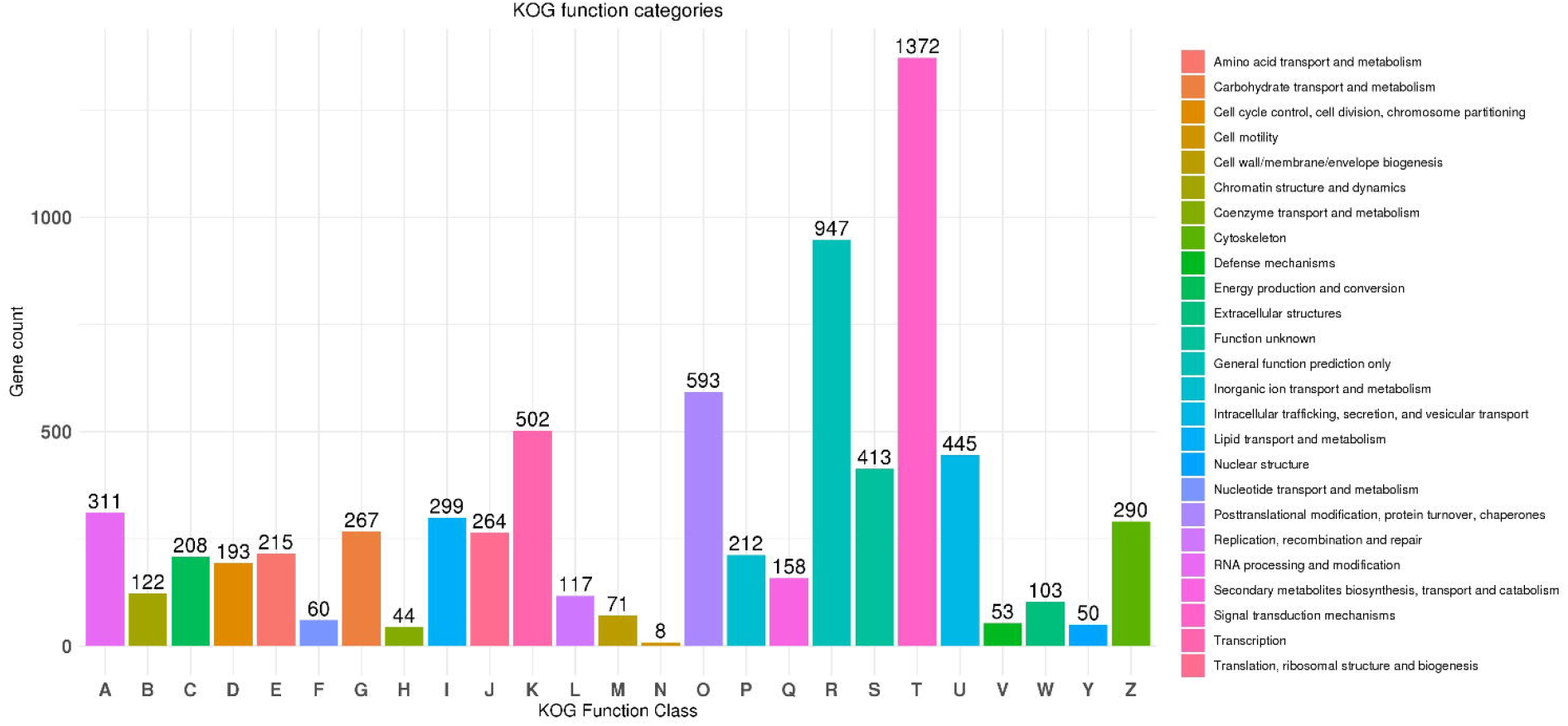
Phylogenetic tree analysis of candidate NPC1 gene from *O. longicollis*. The coleopteran insects are considered as ingroup [*Dendroctonus ponderosae* (Dp: 3 sequences) *Asbolus verrucosus* (Av: 2 sequences), *Tribolium castaneum* (Tc: 3 sequences)] and others are considered as out group [*Drosophila melanogaster* (Dm: 3 sequences); *Rhodnius prolixus* (Rp: 2 sequences). The motif protein sequences only from above insects are used for phylogenetic tree construction. The tree was constructed using PhyML (Guindon et al., 2010) under the best model of WAG +G+I [WAG (Whelan and Goldman, 2001) + Gamma rates (G) + Invariant sites (I)] models of substitution selected by PhyML-SMS (Lef or t et al., 2017), and is based on the amino acid alignment using mafft (Katoh and Standley, 2013). Bootstrap values were estimated using an approximate likelihood ratio test and are displayed as circles on the branches, and are proportionate to the circle diameter (0–500). NPC1- *O. longicollis* sequence was used in this analysis are listed in Supplementary file 1.

### Identification of odorant binding proteins

We identified 19 transcripts encoding putative OBPs in female *O. longicollis* antennal transcriptomes. Among these genes, 16 had complete open reading frames (ORFs) ≥ 400 bp with signal peptides. OBPs commonly called as solubilizers and carriers of odorants and sex pheromones, facilitating the transport of odorant molecules through the sensillar lymph and serving as the liaison between the external environment and olfactory system (Leal 2013; Liu *et al*. 2014). Odorant binding proteins typically has 120 to 150 amino acids and a mass of approximately 14 kDa, including a signal peptide. Many researchers have shown that insect OBPs are found specifically in antennae and other tissues (Gong *et al*. 2009). Similarly, the presence of OBPs in antennae of *O. longicollis* suggesting that their putative role in the odorant detection.

### Identification of Olfactory Receptor and Co-receptors

We have identified 6 putative Odorant Receptors by analysing the female antennal transcriptome. The Simple Modular Architecture Research Tool (SMART) protein domain annotation tool showed that OlOr-2 had seven transmembrane domains, and 4 ORs (OlOr-1, OlOr-3, OlOr-4, OlOr-6) lost their transmembrane domains regions from number three to six and OlOr-5 do not have any domain. ORs are larger multigene family encoding seven transmembrane domain proteins and are the key players in insect olfactory reception (Clyne *et al*. 1999; Vosshall *et al*. 1999; Liu *et al*. 2012). More than fifty ORs were reported from many coleopteran insects but we identified only 6 genes, this indicates that further depth sequencing methods may be address the differences in the identification levels of transcripts. These odorant receptors expressed in the female antennae appear to be involved in female specific behaviours such as finding host plants for oviposition and egg laying.

### Identification of predicted sensory neuron membrane proteins

Two transcripts of SNMP1 and one transcript of SNMP2 were detected from the present study. Among, SNMP1, there were Two Ol SNMP 1 a and b found. They are transmembrane and olfactory-specific proteins that are homologous with human CD36 receptors (Rogers *et al*. 2001; Vogt 2009). These SNMP1 subfamily proteins play a major role on pheromone detection (Liu *et al*. 2014).

### Identification of Ionotropic glutamate receptors

The transcriptome assembly analysis led to the identification of 15 IRs. Particularly 3 sequences (OlGluR1, OlGluR6 and OlGluR7) out of 15 sequences had a complete ORF. Eight IRs (OlGluR3, 4, 5, 8 and 9, OlIR75b, OlIR25a3, OlIR-ANF1) had no transmembrane domain. Ionotropic receptor is another chemosensory receptor family for the detection of a variety of chemical molecules that has been characterized in *D. melanogaster* (Benton *et al*. 2009). IRs are a conserved family and have two major groups: the conserved antennal IRs involved in olfaction and the species-specific divergent IRs are expressed in other tissues like gustatory organs (Hussain *et al*. 2006; Croset *et al*. 2010; Ai *et al*. 2013). Some IR involved in a unique function i.e., IR40a on sensing insect repellent DEET (Kain *et al*. 2013), IR64a on acid detection (Ai *et al*. 2010) and IR76b on low-salt detection (Zhang *et al*. 2013). In the present study, we identified five IR25 subtypes (OlIR25 a1, 2, 3 and OliGluR 6 and 7) which may be involved in any one of the above processes in *O. longicollis*.

## Conclusion

This is the first report on identification of chemosensory genes (NPC1 & 2, OBP, OR, IR and SNMP) from antennal transcriptomics of female *O. longicollis*. The entire identified putative chemosensory genes may be involved in host-plant selection and egg-laying preference of female *O. longicollis*. Phylogenetic analyses of NPC concluded that *O. longicollis* proteins are conserved among the coleopteran insects. Further, the comparative analysis of these female specific genes with male antenna of *O. longicollis* will be much helpful to identify the potential target to develop safer pest control strategies through CRISPR-Cas 9 or RNA interference technology in the future.

## List of Abbreviations

BPW: Banana Pseudostem Weevil;
IPM: Integrated Pest Management;
GC: guanine-cytosine;
NPC: Niemann–Pick type C;
OBP: Odorant Binding Protein;
OR: Olfactory receptor;
IR: Ionotropic Receptor;
SNMP: Sensory Neuron Membrane Protein;
DEET: N, N-Diethyl-meta-toluamide

## Declarations

### Ethics approval and consent to participate

Not applicable.

### Consent for publication

Not applicable.

### Availability of data and materia

On request

### Authors’ contributions

MK and BP designed the experiments; MK wrote the manuscript, MK and VS analysed the transcriptomics data and BP and VS edited and reviewed the manuscript. All the authors read and approved the final version of the manuscript

## Legends for figure

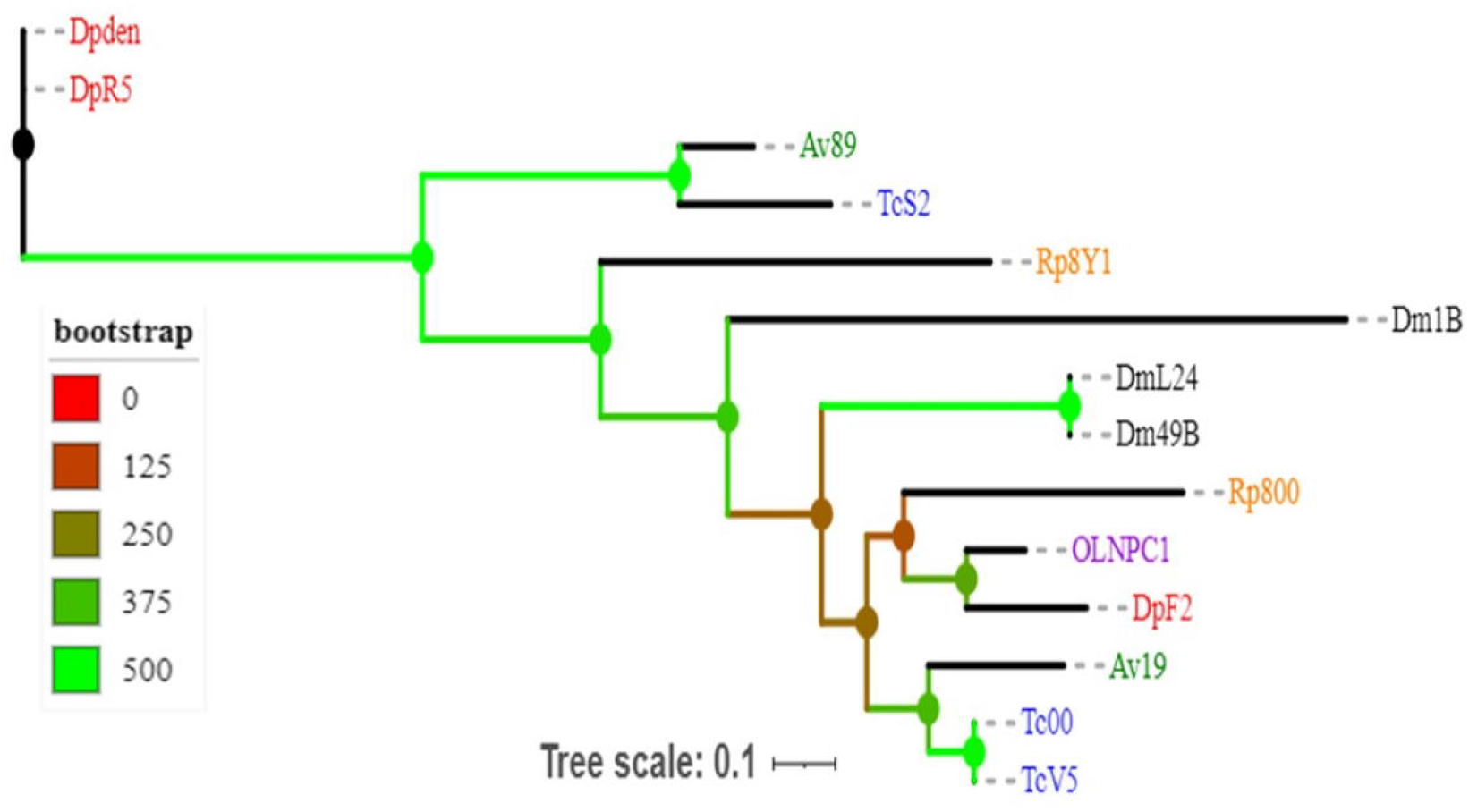

**Table 1.**
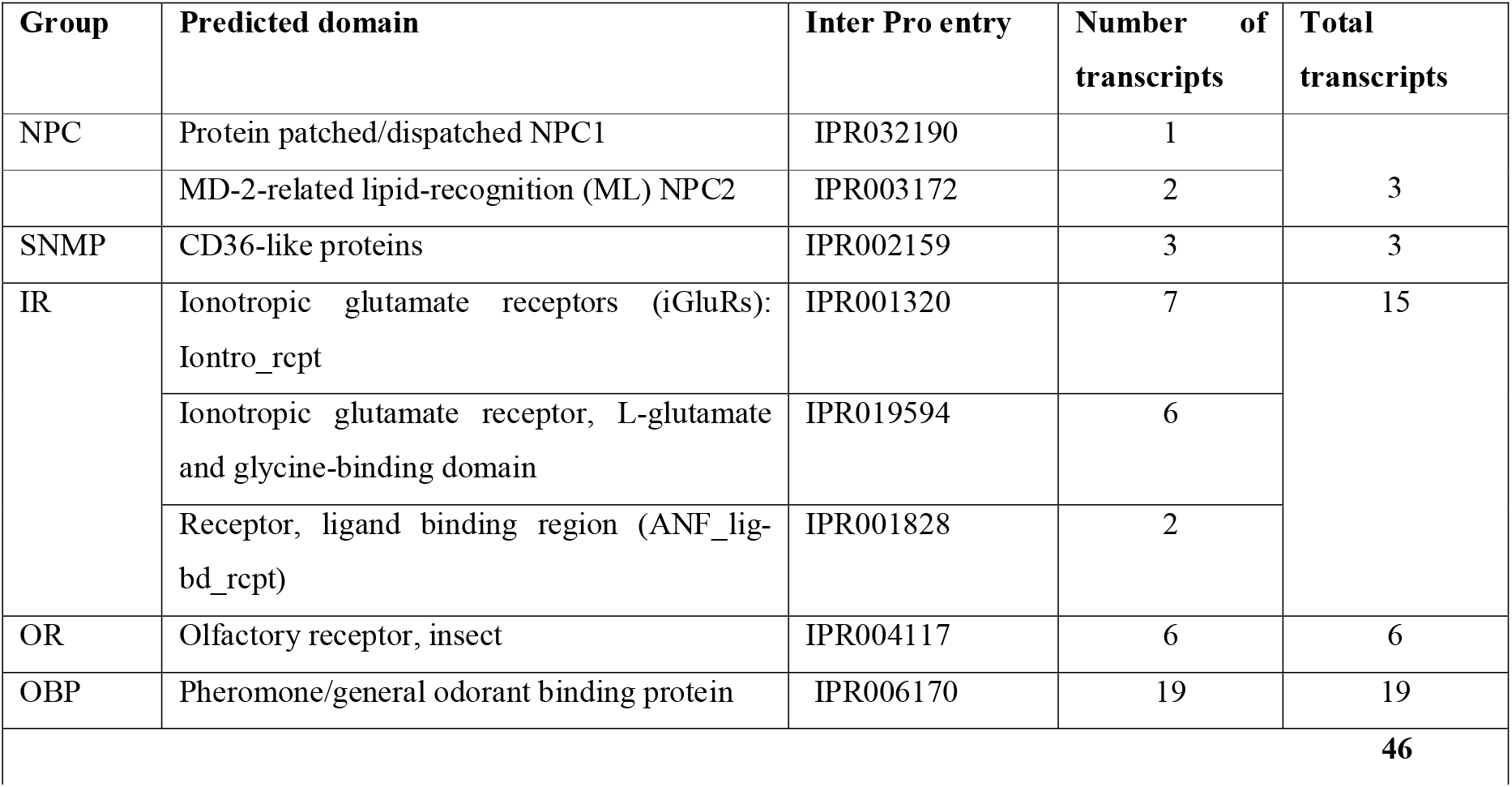
Groups of chemosensory genes, number of transcripts, and their predicted domain with Inter Pro entry according to Blast2GO analysis.

